# Structure of the Arabidopsis receptor kinase SRF6 ectodomain determined from crystals obtained using the LRR crystallisation screen

**DOI:** 10.64898/2026.03.20.713188

**Authors:** Alberto Caregnato, Ulrich Hohmann, Michael Hothorn

**Author notes:** Institute of Molecular Biology, 55128 Mainz, Germany (Ulrich Hohmann).

## Abstract

Plant-specific membrane receptor kinases with structurally diverse extracellular domains regulate key processes in plant growth, development, immunity and symbiosis. Structural studies of these glycoproteins are often hampered by the limited quantities in which they can be obtained. Here, we describe the LRR crystallization screen, which has enabled the successful crystallization and structure determination of multiple receptor kinase ectodomains, including ligand-and co-receptor-bound complexes. As an example, we report the 1.5 Å resolution crystal structure of the leucine-rich repeat (LRR) domain of STRUBBELIG-RECEPTOR FAMILY 6 (SRF6) from *Arabidopsis thaliana*. The SRF6 ectodomain contains seven LRRs and a disulfide-bond-stabilised N-terminal capping domain but lacks the canonical C-terminal cap and the N-glycosylation pattern typically observed in other family members. Previously reported protein-protein interactions between the SRF6 and SRF7 ectodomains and the receptor kinases BRI1, BRL1, BRL3, SERK3 and BIR1-3 could not be confirmed by quantitative isothermal titration calorimetry and grating-coupled interferometry assays, suggesting that these structurally conserved LRR receptor kinases may have signalling functions outside the brassinosteroid pathway.

**Synopsis:** A crystallisation screen that has enabled the structural analysis of various extracellular domains of plant membrane receptor kinases is described together.

## 1. Introduction

Plant genomes harbour a large family of unique membrane receptor kinases (RKs) with extracellular leucine-rich repeat (LRR) extracellular domains (Shiu & Bleecker, 2001; Zhang *et al*., 2026). Members of this family include receptor kinases that bind small molecule, peptide or protein ligands (Hohmann *et al*., 2017; Zhang *et al*., 2017; Moussu & Santiago, 2019), co-receptor kinases required for receptor activation (Brandt & Hothorn, 2016), and receptor pseudo-kinases that regulate the formation of active signalling complexes (Ma *et al*., 2017; Hohmann, Nicolet *et al*., 2018). The extracellular LRR domains of plant RKs are often stabilised by disulphide bridges and are extensively N-glycosylated (Di Matteo *et al*., 2003; Hothorn *et al*., 2011; She *et al*., 2011; Jia *et al*., 2024). Consequently, crystallography studies on plant LRR-RK largely rely on ectodomains obtained by secreted expression in baculovirus-infected insect cells (Hothorn *et al*., 2011; She *et al*., 2011). The resulting low protein yields, often further reduced by enzymatic de-glycosylation prior to crystallisation (Okuda *et al*., 2020), can severely limit the amount of sample available for high-throughput crystallisation screening.

Testing a large number of variables that may influence sample crystallisation (McPherson, 2004) often relies on commercial crystallisation screens, covering conditions previously associated with crystallisation success (Berry *et al*., 2006), or derived from systematic approaches (Gorrec, 2015; Gorrec & Bellini, 2022). Here we report the LRR crystallisation screen, which has yielded a relatively large number of plant LRR-RK ectodomain structures with minimal sample requirements. As an example, we describe the crystallisation and structure solution of the LRR-RK STRUBBELIG-RECEPTOR FAMILY 6 (SRF6).

SRF6 is part of a small protein family of plant receptor kinases. Its founding member STRUBBELIG (SUB/SRF9) was originally described in a genetic screen for mutants defective in ovule development (Schneitz *et al*., 1997) and is involved in plant organ development and cell wall signalling (Chevalier *et al*., 2005; Kwak & Schiefelbein, 2007; Kwak *et al*., 2005; Eyüboglu *et al*., 2007; Chaudhary *et al*., 2020, 2021). The kinase activity of SUB appears to be dispensable for signalling (Chevalier *et al*., 2005). Interaction of SUB with other membrane-integral proteins has been reported (Fulton *et al*., 2009; Vaddepalli *et al*., 2014; Chen *et al*., 2023), but a validated ligand or interaction partner for its extracellular LRR domain remains to be identified.

Mutations in the STRUBBELIG-RECEPTOR FAMILY members SRF3 result in altered immune responses (Alcázar *et al*., 2010, 2014; Atanasov *et al*., 2018; Duan *et al*., 2024) and iron homeostasis (Platre *et al*., 2022). For SRF6, functions in brassinosteroid signalling (Eyüboglu *et al*., 2007; Smakowska-Luzan *et al*., 2018) and in the perception of the cell wall breakdown product trigalacturonic acid (Bhasin *et al*., 2025) have been proposed. Here we report the crystal structure of the SRF6 ectodomain refined at 1.5 Å resolution, and biochemically dissect its potential role in brassinosteroid hormone signalling (Nolan *et al*., 2020), which is mediated by the steroid receptor LRR-RKs BRI1, BRL1 and BRL3 (Li & Chory, 1997; Caño-Delgado *et al*., 2004; Caregnato *et al*., 2025), the LRR co-receptor kinases SERK1-4 (Li *et al*., 2002; Nam & Li, 2002; Gou *et al*., 2012; Hohmann, Santiago *et al*., 2018) and the LRR receptor pseudokinases BIR1-4 (Imkampe *et al*., 2017; Hohmann, Nicolet *et al*., 2018).

## 2. Material and methods

### 2.1. Protein expression and purification

The coding sequence of the AtSRF6 LRR ectodomain (residues 26-287, UNIPROT-ID A8MQH3, https://uniprot.org) was obtained as a synthetic gene codon-optimised for expression in *S. frugiperda* (Twist Bioscience), and introduced by Gibson-assembly cloning (Gibson *et al*., 2009) into a modified pFastBac vector (Geneva Biotech), which provides a *D. melanogaster* Bip signal peptide (MKLCILLAVVAFVGLSLD) (Soejima *et al*., 2013), and a *tobacco etch virus* protease (TEV)-cleavable C-terminal StrepII-9xHis tag. For protein expression, *T. ni* (strain Tnao38) (Hashimoto *et al*., 2010) cells were infected with 10 ml of virus in 250 ml of cells at a density of 2.0 x 10^6^ cells ml^-1^, incubated for 24 h at 28 °C and 110 rev min^-1^ and then for another 48 h at 22 °C and 110 rev min^-1^. The secreted SRF6 ectodomain was purified from the supernatant by sequential Ni^2+^ (HisTrap excel; GE Healthcare; equilibrated in 25 mM KP_i_ pH 7.8, 500 mM NaCl) and StrepII (Strep-Tactin XT; IBA; equilibrated in 25 mM Tris pH 8.0, 250 mM NaCl, 1 mM EDTA) affinity chromatography, followed by size-exclusion chromatography on a HiLoad 16/600 Superdex 200pg column (GE Healthcare), equilibrated in 10 mM sodium citrate pH 5.0, 250 mM NaCl. The monomeric peak fraction was concentrated to 14 mg ml^-1^ using an Amicon Ultra concentrator (molecular-weight cutoff 10,000; Millipore) and directly used for protein crystallisation.

The protein samples used to assess the LRR screen were purified as previously described: SERK1 (12 mg ml^-1^ in 25 mM citric acid, pH 5.0, 100 mM NaCl) (Santiago *et al*., 2013), BRI1 – BLD – SERK1 (10 mg ml^-1^ in 25 mM citric acid, pH 5.0, 100 mM NaCl) (Santiago *et al*., 2013), HAESA – IDA (5 mg ml^-1^ in 20 mM citric acid pH 5.0, 100 mM NaCl) (Santiago *et al*., 2016), HAESA – IDA – SERK1 (12 mg ml^-1^ in 20 mM citric acid pH 5.0, 100 mM NaCl) (Santiago *et al*., 2016), BIR2 (9 mg ml^-1^ in 20 mM sodium citrate pH 5.0, 150 mM NaCl) (Hohmann, Nicolet *et al*., 2018), BIR3 – SERK1 (14 mg ml^-1^ in 20 mM sodium citrate pH 5.0, 150 mM NaCl) (Hohmann, Nicolet *et al*., 2018), PDLP5 (70 mg ml^-1^ in 20 mM sodium citrate pH 5.0, 150 mM NaCl) (Vaattovaara *et al*., 2019), SOBIR1 (20 mg ml^-1^ in 20 mM sodium citrate pH 5.0, 150 mM NaCl) (Hohmann & Hothorn, 2019), GSO1 – CIF2 (1 mg ml^-1^ in 20 mM sodium citrate pH 5.0, 150 mM NaCl) (Okuda *et al*., 2020), BRI1 – A1JME (6 mg ml^-1^ in 20 mM citric acid pH 5.0, 250 mM NaCl) (Caregnato *et al*., 2025), BRL3 – A1JMF (7 mg ml^-1^ in 20 mM citric acid pH 5.0, 250 mM NaCl) (Caregnato *et al*., 2025), and BRL2 (6 mg ml^-1^ in 20 mM citric acid pH 5.0, 250 mM NaCl) (Caregnato *et al*., 2025).

For isothermal titration calorimetry (ITC) and grating-coupled interferometry (GCI) assays AtSRF7 (residues 26-287, UNIPROT-ID B5X583) was cloned, expressed and purified as described for SRF6. BRI1, BRL1, BRL3 and SERK3 were expressed and purified as described previously (Hohmann, Santiago *et al*., 2018; Caregnato *et al*., 2025), as was the expression and purification of BIR1-4 (Hohmann, Nicolet *et al*., 2018). For ITC assays, the C-terminal affinity tags in BRL1 and SRF7 were removed by TEV cleavage over night at 4 °C, followed by size-exclusion chromatography into ITC buffer (25 mM sodium citrate pH 5.0, 150 mM NaCl). BRI1, BRL1 and BRL3 biotinylation for grating-coupled interferometry was achieved by incubating the respective Avi-tagged receptor with His-tagged BirA (Fairhead & Howarth, 2015). The receptor at a final concentration of 20 µM was incubated for 1 h at 30 °C with BirA, biotin, ATP and MgCl_2_ at final concentrations of 85 μM, 150 μM, 2 mM and 5 mM respectively. The BirA enzyme was subsequently removed by Ni^2+^ affinity chromatography, and the biotinylated receptor was concentrated and loaded onto a HiLoad Superdex 200 16/200 pg column (Cytiva) equilibrated in 20 mM sodium citrate pH 5.0, 150 mM NaCl. The C-terminal affinity tags from SRF6, SRF7, SERK3 and BIR1-4 were removed for these assays, as described above.

### 2.2. LRR protein crystallisation screening

The LRR crystallisation screen was prepared in 96 50 ml falcon tubes from stock solutions. Precipitants: PEG 1,000 (Sigma #81188, 50 % [w/v] stock solution), PEG 3,350 (Sigma #88276, 50 % [w/v] stock solution), PEG 8,000 (Sigma #89510, 40% [w/v] stock solution), sodium malonate (3.4 M stock solution, pH 4.0-8.0, Hampton Research), ammonium sulfate (Sigma #A4418, 4 M stock solution). Salts: ammonium sulfate (Sigma #A4418, 2 M stock solution), lithium sulfate (Sigma #13029, 0.5 M stock solution), sodium chloride (Sigma #S9888, 2 M stock solution), sodium citrate (0.5 M stock solution from 0.5 M citric acid [#Sigma 251275] with 1.5 M sodium hydroxide), ammonium acetate (Sigma #09688, 0.5 M stock solution), magnesium chloride (Sigma #M2393, 0.5 M stock solution), lithium chloride (Sigma #L9650, 2 M stock solution). Buffers: citric acid/NaOH pH 4.0 (Sigma #251275, 1 M stock solution), sodium acetate/acetic acid pH 5.5 (Sigma #S8750, 1 M stock solution), Bis-Tris/HCl pH 7.0 (Sigma B7535, 1 M stock solution), Tris base/HCl pH 8.5 (Sigma #T4661, 1 M stock solution). Crystallisation experiments were performed at room temperature in 96 well MRC 2 well sitting drop plates (SWISSCI #MRC96T-UVP). Drops were composed of 0.2 μl of protein solution and 0.2 μl of crystallisation buffer suspended over 100 μl of the latter as reservoir solution. The second drop was set up with the protein diluted 1:3 in protein storage buffer. Plates were inspected after 24 h, 3 d and 2 month on a CX31 microscope (Olympus). In the case of AtSRF6, diffraction quality crystals appeared after 3 d in LRR screen conditions C5 (25% [w/v] PEG 3,350, 0.2 M sodium chloride, 0.1 M citric acid, pH 4.0) and F9 (20% [w/v] PEG 8,000, 0.2 M magnesium chloride, 0.1 M citric acid, pH 4.0). A needle-shaped crystal (∼300 x 50 x 50 μm) from condition F9 was transferred to reservoir solution supplemented with 15% (v/v) glycerol and snap-frozen in liquid N_2_.

### 2.3. Crystallographic data collection, structure solution and refinement

Redundant sulphur single-wavelength anomalous dispersion (SAD) data (λ= 2.079 Å, five 360° wedges at 0.1° oscillation, with Χ set to -20°, -10°, 0°, 10°, 20°) to 2.3 Å were collected at beam line X06DA of the Swiss Light Source (SLS), Villigen, Switzerland equipped with a Pilatus 2M-F detector (Dectris Ltd.) and a multi-axis goniometer. Next, the needle-shaped crystal was translated and an additional high-resolution native dataset (λ= 0.978 Å, one 360° wedge at 0.1° oscillation) was collected to 1.5 Å resolution. Data were processed and scaled with XDS and XSCALE, respectively (Kabsch, 1993). Analysis with phenix.xtriage (Zwart *et al*., 2005) indicated that the anomalous signal of the scaled SAD dataset extended only to ∼ 4.5 Å. Therefore, the native dataset was input into the MORDA automatic molecular replacement (MR) pipeline (https://www.ccp4.ac.uk/morda-automatic-molecular-replacement-pipeline/), which returned a marginal solution in space-group *P*4_3_2_1_2 (translation function Z-score 8.4, R_work_/R_free_ 0.486/0.495) with a single molecule of the previously reported Arabidopsis POLLEN RECEPTOR-LIKE KINASE 6 (PRK6) ectodomain structure in the asymmetric unit (PDB-ID: pdb_00005y9w) (Zhang *et al*., 2017). The MORDA solution was input into Phaser for MR-SAD phasing against the SAD dataset at 2.3 Å (starting figure of merit [FOM] was 0.309 at 2.3 Å resolution), yielding four putative sulphur sites by log-likelihood-gradient completion (final FOM was 0.475). After merging with the high-resolution native dataset, these starting phases were used for automatic model building in ARP/wARP (Langer *et al*., 2008). The resulting structure was completed in iterative cycles of manual model correction in COOT (Emsley & Cowtan, 2004) and restrained refinement in REFMAC5 (Murshudov *et al*., 1997). The final model had excellent stereochemistry, assessed with phenix.molprobity (Davis *et al*., 2007). Structural diagrams were prepared with Pymol (https://pymol.org) and ChimeraX (Meng *et al*., 2023). Phased anomalous difference maps were generated with phenix.find_peaks_holes and displayed in ChimeraX.

### 2.4. Analytical size-exclusion chromatography

Analytical size exclusion chromatography (SEC) experiments were performed on a Superdex 200 increase 10/300 GL column (GE Healthcare) pre-equilibrated in 20 mM sodium citrate pH 5.0, 250 mM NaCl. 500 μg of a mixture containing the BRL1 and SRF6 ectodomains in 1:1 molar ratio were injected in a volume of 100 μl onto the column and elution at 0.75 ml min^-1^ was monitored by ultraviolet absorbance at λ = 280 nm. Peak fractions were analysed by SDS-PAGE.

### 2.5. Isothermal titration calorimetry

The ITC experiment was performed at 25 °C using a Nano ITC (TA Instruments) with a 1.0 ml standard cell and a 250 μl titration syringe. SRF6 and BRL1 were gel filtrated into ITC buffer (25 mM sodium citrate pH 5.0, 150 mM NaCl). 10 μl SRF7 aliquots (∼220 μM) were injected into into ∼23 μM BRL1 in the cell at 150-s intervals. ITC data were corrected for the heat of dilution by subtracting the mixing enthalpies for titrant solution injections into protein-free ITC buffer. Data were analysed using the NanoAnalyze program (version 3.5) as provided by the manufacturer.

### 2.6. SRF-family phylogeny

A multiple sequence alignment of the SRF1 (TAIR-ID: AT2G20850, https://www.arabidopsis.org/), SRF2 (AT5G06820), SRF3 (AT4G03390), SRF5 (AT1G78980), SRF6 (AT1G53730), SRF7 (AT3G14350), SRF8 (AT4G22130) and SRF9/SUB (AT1G11130) protein sequences was generated with probalign (Roshan & Livesay, 2006). The phylogenetic tree was generated with iqtree2 (Minh *et al*., 2020) and displayed in figtree (http://tree.bio.ed.ac.uk/software/figtree/).

### 2.7. Grating-coupled interferometry

GCI assays were performed on a Creoptix WAVE system (Malvern Panalytical). Binding of the isolated SRF6, SRF7 and SERK3 (positive control) was measured by amine-coupling the BRI1, BRL1, BRL3, BIR1, BIR2, BIR3 or SERK3 ectodomains (ligand) onto 2PCP WAVEchips (quasi-planar polycarboxylate surface; Creoptix AG, Switzerland). Chips were conditioned using borate buffer (100 mM sodium borate pH 9.0, 1 M NaCl; Xantec, Germany), and the respective ligand were immobilized on the chip surface via a standard amine-coupling protocol, which consisted of 7 min activation with a 1:1 mix of 400 mM N-(3-dimethylaminopropyl)-N’-ethylcarbodiimide hydrochloride and 100 mM N-hydroxysuccinimide (both Xantec, Germany), followed by injection of the ligand (1 to 50 μg ml^-1^) in 10 mM sodium acetate pH 5.0 (Sigma, Germany) until the desired density was reached, and quenched with 1 M ethanolamine pH 8.0 for 7 min (Xantec, Germany). BSA (0.5% in 10 mM sodium acetate pH 5.0; BSA from Roche, Switzerland) was used to passivate the surface between ligand injection and ethanolamine quenching. The isolated SRF6, SRF7 or SERK3 ectodomains were used as analyte. Kinetic analyses were performed at 25°C with a 1:2 dilution series from 2 μM in 20 mM sodium citrate pH 5.0, 250 mM NaCl, 100 nM brassinolide, with blank injections for double referencing and DMSO calibration for bulk correction. Data correction and analysis was performed with the Creoptix WAVEcontrol software (corrections applied: X and Y offset, DMSO calibration and double referencing). Data were fitted to either one-to-one binding models or mass-transport limited models, using bulk correction.

## 3. Results

### 3.1. A high-throughput crystallisation screen for extracellular LRR proteins

The recombinant expression and purification of the first plant LRR-RK ectodomain in baculovirus-infected insect cells yielded only 50–200 µg of purified BRI1 per litre of cell culture (Hothorn *et al*., 2011). For the subsequent crystallisation of the BRI1 – BLD – SERK1 complex (Santiago *et al*., 2013), a tailored crystallisation screen was therefore developed. The screen simply combined the crystallisation conditions reported for the few LRR domain structures that had been previously reported for different animal and plant extracellular proteins (Table 1) (Uff *et al*., 2002; He *et al*., 2003; Di Matteo *et al*., 2003; Kim *et al*., 2005; Choe *et al*., 2005; Bell *et al*., 2005; Liu *et al*., 2008; Han *et al*., 2008; Hothorn *et al*., 2011; She *et al*., 2011). The conditions were replicated using a buffer system covering a pH range of 4.0–8.5 (the mean pH of the screen is 5.4; Table 2). This was based on the idea that extracellular proteins usually reside in an acidic environment (the pH range of the plant apoplast is 4.5-6.5) (Almeida & Huber, 1999) and that the mean pH of many commercially available, high-throughput crystallisation screens was close to neutral (Crystal Screen HT: pH 6.6; Index HT: mean pH 6.8; PEG Ion: pH 6.8; Hampton Research).

**Table 1.** LRR ectodomain crystallisation conditions reported between 2002 – 2011.

| Protein | Precipitant | Salt | Buffer | Reference |
| --- | --- | --- | --- | --- |
| GPIba | 2 M ammonium sulfate |  | 0.1 M Tris/HCl pH 6.5 | (Uff *et al.*, 2002) |
| NgR | 35% PEG 4,000 | 0.25 M sodium chloride | 0.1 M sodium acetate pH 6.5 | (He *et al.*, 2003) |
| PGIP | 20-30% PEG 4,000 | 0.2 M ammonium acetate | 0.1 M sodium acetate pH 4.6 | (Di Matteo *et al.*, 2003) |
| CD14 | 1.9 M lithium sulfate | 0.005 M nickel chloride | 0.1 M Hepes pH 7.5 | (Kim *et al.*, 2005) |
| TLR3 | 23% PEG 2,000 | 0.4 M ammonium sulfate |  | (Choe *et al.*, 2005) |
| TL3 | 25% PEG 4,000 | 0.2 M lithium sulfate | 0.1 M sodium acetate pH 4.6 | (Bell *et al.*, 2005) |
| mTL3 | 15% PEG,6000 / 5% Dextran 5,000 | 1 M lithium chloride | 0.1 M citric acid pH 4.0 | (Liu *et al.*, 2008) |
| mTLR3 - dsRNA | 13% PEG 3,350 / 5% Dextran 5,000 | 0.2 M ammonium citrate pH 5.5 |  | (Liu *et al.*, 2008) |
| VLR - RBC36 | 2 M ammonium sulfate | 0.2 M sodium / potassium tartrate | 0.1 M sodium citrate pH 5.6 | (Han *et al.*, 2008) |
| BRI1 – BLD | 14% PEG 4,000 | 0.2 M ammonium sulfate | 0.1 M citric acid pH 4.0 | (Hothorn *et al.*, 2011) |
| BRI1 – BLD | 20% PEG 3,350 | 0.2 M ammonium sulfate |  | (She *et al.*, 2011) |

**Table 2.** LRR screen formulation.

| Well | Precipitant | Salt | Buffer |
| --- | --- | --- | --- |
| A1 | 30% PEG 1,000 | 0.2 M ammonium sulfate | 0.1 M citric acid pH 4.0 |
| A2 | 30% PEG 1,000 | 0.2 M ammonium sulfate | 0.1 M sodium acetate pH 5.5 |
| A3 | 30% PEG 1,000 | 0.2 M ammonium sulfate | 0.1 M Bis-Tris pH 7.0 |
| A4 | 30% PEG 1,000 | 0.2 M ammonium sulfate | 0.1 M Tris pH 8.5 |
| A5 | 25% PEG 3,350 | 0.2 M ammonium sulfate | 0.1 M citric acid pH 4.0 |
| A6 | 25% PEG 3,350 | 0.2 M ammonium sulfate | 0.1 M sodium acetate pH 5.5 |
| A7 | 25% PEG 3,350 | 0.2 M ammonium sulfate | 0.1 M Bis-Tris pH 7.0 |
| A8 | 25% PEG 3,350 | 0.2 M ammonium sulfate | 0.1 M Tris pH 8.5 |
| A9 | 20% PEG 8,000 | 0.2 M ammonium sulfate | 0.1 M citric acid pH 4.0 |
| A10 | 20% PEG 8,000 | 0.2 M ammonium sulfate | 0.1 M sodium acetate pH 5.5 |
| A11 | 20% PEG 8,000 | 0.2 M ammonium sulfate | 0.1 M Bis-Tris pH 7.0 |
| A12 | 20% PEG 8,000 | 0.2 M ammonium sulfate | 0.1 M Tris pH 8.5 |
| B1 | 30% PEG 1,000 | 0.2 M lithium sulfate | 0.1 M citric acid pH 4.0 |
| B2 | 30% PEG 1,000 | 0.2 M lithium sulfate | 0.1 M sodium acetate pH 5.5 |
| B3 | 30% PEG 1,000 | 0.2 M lithium sulfate | 0.1 M Bis-Tris pH 7.0 |
| B4 | 30% PEG 1,000 | 0.2 M lithium sulfate | 0.1 M Tris pH 8.5 |
| B5 | 25% PEG 3,350 | 0.2 M lithium sulfate | 0.1 M citric acid pH 4.0 |
| B6 | 25% PEG 3,350 | 0.2 M lithium sulfate | 0.1 M sodium acetate pH 5.5 |
| B7 | 25% PEG 3,350 | 0.2 M lithium sulfate | 0.1 M Bis-Tris pH 7.0 |
| B8 | 25% PEG 3,350 | 0.2 M lithium sulfate | 0.1 M Tris pH 8.5 |
| B9 | 20% PEG 8,000 | 0.2 M lithium sulfate | 0.1 M citric acid pH 4.0 |
| B10 | 20% PEG 8,000 | 0.2 M lithium sulfate | 0.1 M sodium acetate pH 5.5 |
| B11 | 20% PEG 8,000 | 0.2 M lithium sulfate | 0.1 M Bis-Tris pH 7.0 |
| B12 | 20% PEG 8,000 | 0.2 M lithium sulfate | 0.1 M Tris pH 8.5 |
| C1 | 30% PEG 1,000 | 0.2 M sodium chloride | 0.1 M citric acid pH 4.0 |
| C2 | 30% PEG 1,000 | 0.2 M sodium chloride | 0.1 M sodium acetate pH 5.5 |
| C3 | 30% PEG 1,000 | 0.2 M sodium chloride | 0.1 M Bis-Tris pH 7.0 |
| C4 | 30% PEG 1,000 | 0.2 M sodium chloride | 0.1 M Tris pH 8.5 |
| C5 | 25% PEG 3,350 | 0.2 M sodium chloride | 0.1 M citric acid pH 4.0 |
| C6 | 25% PEG 3,350 | 0.2 M sodium chloride | 0.1 M sodium acetate pH 5.5 |
| C7 | 25% PEG 3,350 | 0.2 M sodium chloride | 0.1 M Bis-Tris pH 7.0 |
| C8 | 25% PEG 3350 | 0.2 M sodium chloride | 0.1 M Tris pH 8.5 |
| C9 | 20% PEG 8,000 | 0.2 M sodium chloride | 0.1 M citric acid pH 4.0 |
| C10 | 20% PEG 8,000 | 0.2 M sodium chloride | 0.1 M sodium acetate pH 5.5 |
| C11 | 20% PEG 8,000 | 0.2 M sodium chloride | 0.1 M Bis-Tris pH 7.0 |
| C12 | 20% PEG 8,000 | 0.2 M sodium chloride | 0.1 M Tris pH 8.5 |
| D1 | 30% PEG 1,000 | 0.2 M sodium citrate | 0.1 M citric acid pH 4.0 |
| D2 | 30% PEG 1000 | 0.2 M sodium citrate | 0.1 M sodium acetate pH 5.5 |
| D3 | 30% PEG 1,000 | 0.2 M sodium citrate | 0.1 M Bis-Tris pH 7.0 |
| D4 | 30% PEG 1,000 | 0.2 M sodium citrate | 0.1 M Tris pH 8.5 |
| D5 | 25% PEG 3,350 | 0.2 M sodium citrate | 0.1 M citric acid pH 4.0 |
| D6 | 25% PEG 3,350 | 0.2 M sodium citrate | 0.1 M sodium acetate pH 5.5 |
| D7 | 25% PEG 3,350 | 0.2 M sodium citrate | 0.1 M Bis-Tris pH 7.0 |
| D8 | 25% PEG 3,350 | 0.2 M sodium citrate | 0.1 M Tris pH 8.5 |
| D9 | 20% PEG 8,000 | 0.2 M sodium citrate | 0.1 M citric acid pH 4.0 |
| D10 | 20% PEG 8,000 | 0.2 M sodium citrate | 0.1 M sodium acetate pH 5.5 |
| D11 | 20% PEG 8,000 | 0.2 M sodium citrate | 0.1 M Bis-Tris pH 7.0 |
| D12 | 20% PEG 8,000 | 0.2 M sodium citrate | 0.1 M Tris pH 8.5 |
| E1 | 30% PEG 1,000 | 0.2 M ammonium acetate | 0.1 M citric acid pH 4.0 |
| E2 | 30% PEG 1,000 | 0.2 M ammonium acetate | 0.1 M sodium acetate pH 5.5 |
| E3 | 30% PEG 1,000 | 0.2 M ammonium acetate | 0.1 M Bis-Tris pH 7.0 |
| E4 | 30% PEG 1,000 | 0.2 M ammonium acetate | 0.1 M Tris pH 8.5 |
| E5 | 25% PEG 3,350 | 0.2 M ammonium acetate | 0.1 M citric acid pH 4.0 |
| E6 | 25% PEG 3,350 | 0.2 M ammonium acetate | 0.1 M sodium acetate pH 5.5 |
| E7 | 25% PEG 3,350 | 0.2 M ammonium acetate | 0.1 M Bis-Tris pH 7.0 |
| E8 | 25% PEG 3,350 | 0.2 M ammonium acetate | 0.1 M Tris pH 8.5 |
| E9 | 20% PEG 8,000 | 0.2 M ammonium acetate | 0.1 M citric acid pH 4.0 |
| E10 | 20% PEG 8,000 | 0.2 M ammonium acetate | 0.1 M sodium acetate pH 5.5 |
| E11 | 20% PEG 8,000 | 0.2 M ammonium acetate | 0.1 M Bis-Tris pH 7.0 |
| E12 | 20% PEG 8,000 | 0.2 M ammonium acetate | 0.1 M Tris pH 8.5 |
| F1 | 30% PEG 1,000 | 0.2 M magnesium chloride | 0.1 M citric acid pH 4.0 |
| F2 | 30% PEG 1,000 | 0.2 M magnesium chloride | 0.1 M sodium acetate pH 5.5 |
| F3 | 30% PEG 1,000 | 0.2 M magnesium chloride | 0.1 M Bis-Tris pH 7.0 |
| F4 | 30% PEG 1,000 | 0.2 M magnesium chloride | 0.1 M Tris pH 8.5 |
| F5 | 25% PEG 3,350 | 0.2 M magnesium chloride | 0.1 M citric acid pH 4.0 |
| F6 | 25% PEG 3,350 | 0.2 M magnesium chloride | 0.1 M sodium acetate pH 5.5 |
| F7 | 25% PEG 3,350 | 0.2 M magnesium chloride | 0.1 M Bis-Tris pH 7.0 |
| F8 | 25% PEG 3,350 | 0.2 M magnesium chloride | 0.1 M Tris pH 8.5 |
| F9 | 20% PEG 8,000 | 0.2 M magnesium chloride | 0.1 M citric acid pH 4.0 |
| F10 | 20% PEG 8,000 | 0.2 M magnesium chloride | 0.1 M sodium acetate pH 5.5 |
| F11 | 20% PEG 8,000 | 0.2 M magnesium chloride | 0.1 M Bis-Tris pH 7.0 |
| F12 | 20% PEG 8000 | 0.2 M magnesium chloride | 0.1 M Tris pH 8.5 |
| G1 | 3.4 M Na malonate pH 4.0 |  |  |
| G2 | 3.4 M Na malonate pH 5.0 |  |  |
| G3 | 3.4 M Na malonate pH 6.0 |  |  |
| G4 | 3.4 M Na malonate pH 8.0 |  |  |
| G5 | 2.5 M Na malonate pH 4.0 |  |  |
| G6 | 2.5 M Na malonate pH 5.0 |  |  |
| G7 | 2.5 M Na malonate pH 6.0 |  |  |
| G8 | 2.5 M Na malonate pH 8.0 |  |  |
| G9 | 1.8 M Na malonate pH 4.0 |  |  |
| G10 | 1.8 M Na malonate pH 5.0 |  |  |
| G11 | 1.8 M Na malonate pH 6.0 |  |  |
| G12 | 1.8 M Na malonate pH 8.0 |  |  |
| H1 | 3.5 M ammonium sulfate |  | 0.1 M citric acid pH 4.0 |
| H2 | 3.5 M ammonium sulfate |  | 0.1 M sodium acetate pH 5.5 |
| H3 | 3.5 M ammonium sulfate |  | 0.1 M Bis-Tris pH 7.0 |
| H4 | 3.5 M ammonium sulfate |  | 0.1 M Tris pH 8.5 |
| H5 | 3.5 M ammonium sulfate |  | 0.1 M citric acid pH 4.0 |
| H6 | 3.5 M ammonium sulfate |  | 0.1 M sodium acetate pH 5.5 |
| H7 | 3.5 M ammonium sulfate |  | 0.1 M Bis-Tris pH 7.0 |
| H8 | 3.5 M ammonium sulfate |  | 0.1 M Tris pH 8.5 |
| H9 | 20% PEG 3,350 | 1.0 M lithium chloride | 0.1 M citric acid pH 4.0 |
| H10 | 20% PEG 3,350 | 1.0 M lithium chloride | 0.1 M sodium acetate pH 5.5 |
| H11 | 20% PEG 3,350 | 1.0 M lithium chloride | 0.1 M Bis-Tris pH 7.0 |
| H12 | 20% PEG 3,350 | 1.0 M lithium chloride | 0.1 M Tris pH 8.5 |

Despite its rather simple design principle, the LRR crystallisation screen enabled the crystallisation and structural analysis of the steroid receptor kinase BRI1^sud1^, the isolated ectodomain of the co-receptor kinase SERK1, and the BRI1^sud1^ – BLD – SERK1 complex (Santiago *et al*., 2013) (Fig. 1). Over the years, our laboratory has used the LRR screen to determine the structures of several additional plant receptor kinase domains, including isolated LRR ectodomains (Hohmann, Nicolet *et al*., 2018; Hohmann, Santiago *et al*., 2018; Hohmann & Hothorn, 2019), receptor – small molecule and receptor – peptide ligand complexes (Santiago *et al*., 2016; Okuda *et al*., 2020; Caregnato *et al*., 2025), as well as receptor – co-receptor or regulatory complexes (Santiago *et al*., 2016; Hohmann, Nicolet *et al*., 2018) (Fig. 1). Outside our laboratory, the screen has been used to crystallise the ectodomains of the cell wall protein LRX4 in complex with its peptide ligand RALF4 (conditions E6, E7) (Moussu *et al*., 2020), the LRR-RK HSL1 (condition A1) (Roman *et al*., 2022), and the related HSL3 (condition E5) (Jiménez-Sandoval *et al*., 2025). Finally, the LRR screen was used to crystallise the plant receptor-like protein PDLP5, which has a non-LRR tandem-malectin ectodomain (Vaattovaara *et al*., 2019) (Fig. 1).

**Figure 1.**
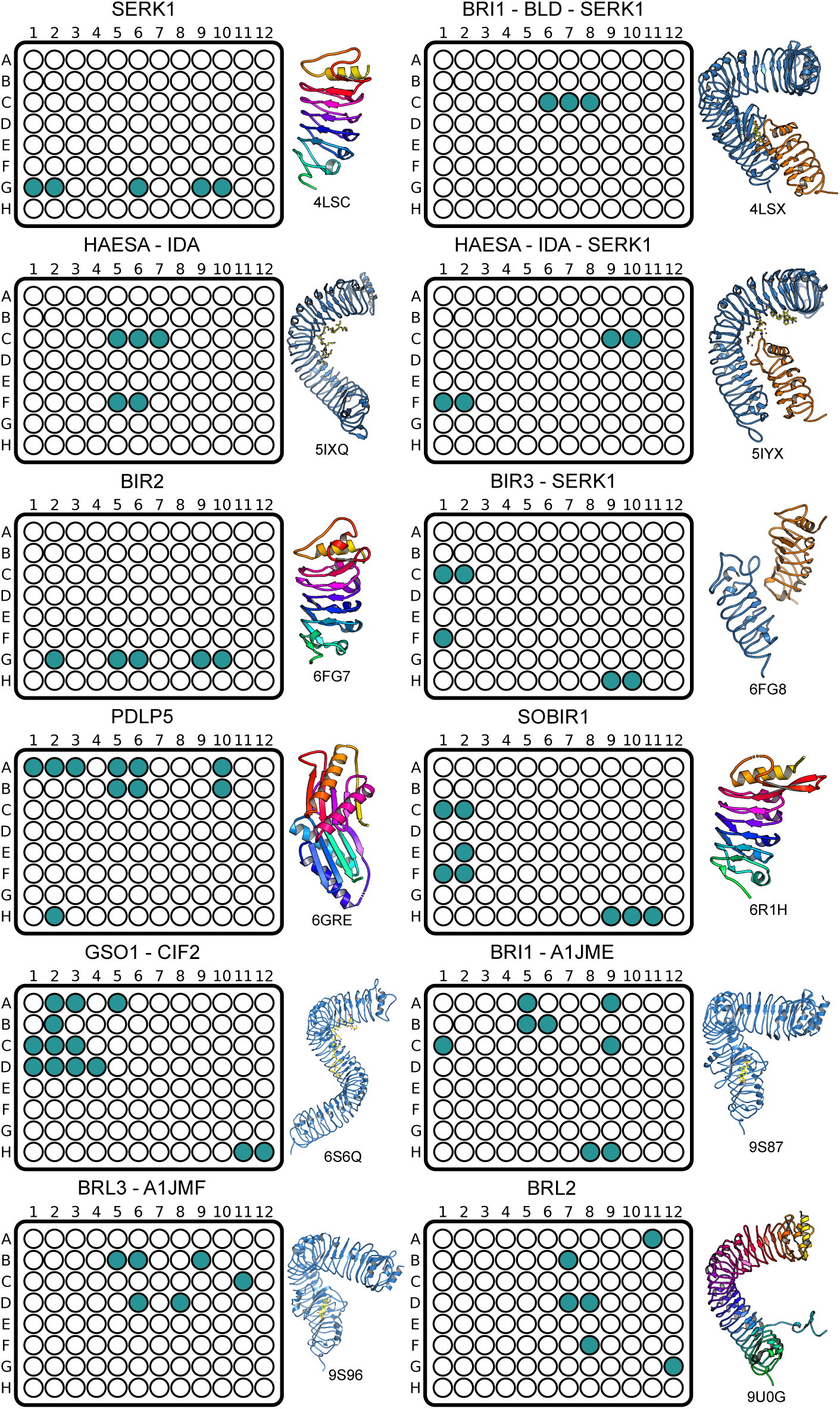
Initial crystallisation hits for 10 different plant receptor kinases obtained using the LRR crystallisation screen. Shown are schematic representations of a 96 well high-throughput crystallisation screen plate with positive hits highlighted in cyan, observed after 2 month of incubation at room temperature. A ribbon diagram of the respective structure is shown alongside, with isolated ectodomain structures coloured from red (N-terminus) to green (C-terminus). Receptor-ligand complexes are shown in blue (ribbon diagram) and yellow (in bonds representations), respectively. Co-receptor kinases are depicted in orange.

### 3.2. Structure solution of SRF6 crystallised with the LRR screen

Next, we used the LRR screen to determine the structure of the isolated LRR ectodomain of the *Arabidopsis thaliana* receptor kinase SRF6 (see Fig. 2a). SRF6 crystallised readily in various conditions in the screen, forming diffraction-quality, needle-shaped crystals in conditions E9 and F6 after three days’ incubation at room temperature (see Methods) (Fig. 2 *b*). Redundant SAD and a high-resolution native dataset data were collected from a single crystal. The structure of SRF6 was solved by the MR-SAD method implemented in PHASER (McCoy *et al*., 2007). The solution contains a single monomer in the asymmetric unit and four sulphur sites corresponding to a disulphide bridge and to Met42 in the N-terminal capping domain, and to Met91 in the LRR core (Fig. 2c). The final model was refined against the high-resolution native dataset at 1.50 Å resolution (Table 3, Fig. 2 *d*). An example region of the final (2F_o_ - F_c_) map is shown in Fig. 2e. The SRF6 structure reveals a compact LRR domain comprising seven LRRs, rather than six as previously suggested for the related SRF9/SUB (Vaddepalli *et al*., 2011) (Fig. 2d). Several genetic missense alleles identified for SRF9/SUB map to the N-terminal capping domain of SRF6, supporting the function of the N-cap in the folding of LRR domains (Truhlar & Komives, 2008) (shown in magenta in Fig. 2d). In fact, the Cys57-Tyr mutation in SRF9/SUB corresponds to the *bri1-5* allele (Cys69-Tyr), which causes the LRR-RK BRI1 to be retained in the endoplasmic reticulum (Hong *et al*., 2008). In contrast to many other LRR-RK ectodomain structures, SRF6 lacks any visible N-glycans, and its C-terminal capping domain is devoid of a stabilizing disulphide bridge, as previously seen for example in the co-receptor kinase SERK1 (Santiago *et al*., 2013) (Fig. 2f). Instead, the C-terminal cap of SRF6 is structurally reminiscent of the situation described for the immune receptor kinase SOBIR1 (Hohmann & Hothorn, 2019) (Fig. 2f). Taken together, SRF6 shares the overall ectodomain structure with other plant LRR receptor and co-receptor kinases (Hohmann *et al*., 2017).

**Figure 2.**
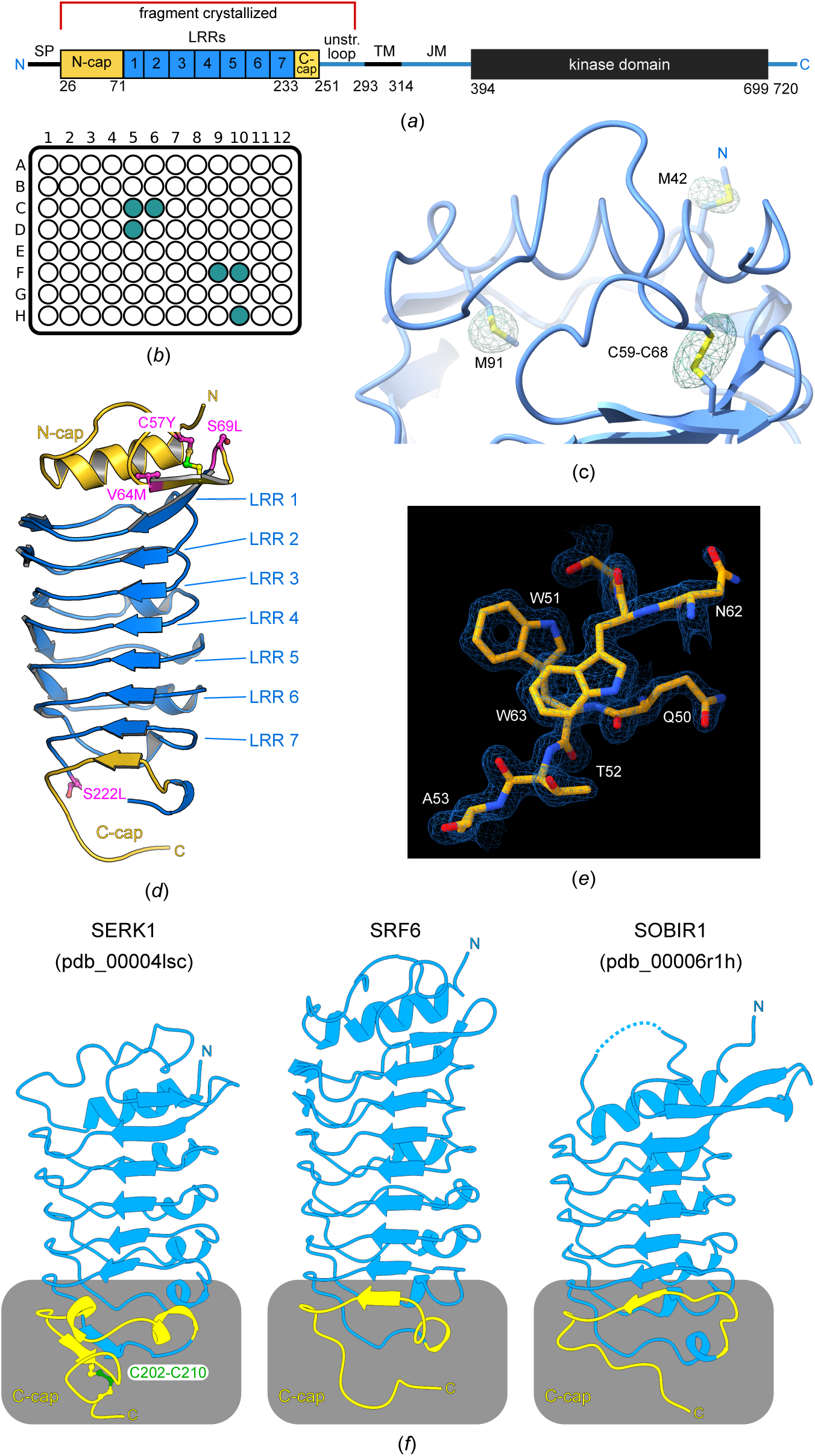
Crystallographic analysis of the LRR ectodomain of the Arabidopsis receptor kinase SRF6. (*a*) Schematic representation of SRF6 (SP, signal peptide; N-cap, N-terminal capping domain; C-cap, C-terminal capping domain; TM, trans-membrane helix; JM, juxta-membrane motif. The crystallised fragment is indicated by a red line. (*b*) Schematic representation of a 96 well high-throughput crystallisation screen plate, conditions containing SRF6 crystals after 3 d of incubation at room temperature are shown in cyan. (*c*) Ribbon diagram of the SRF6 N-terminal capping domain, with Met42, Cys59, Cys68 and Met91 shown in bonds representation and including a phased anomalous difference map contoured at 9 σ (green mesh). (*d*) Overall structure of the SRF6 ectodomain. Shown is a ribbon diagram, the secondary structure was assigned using DSSP (Kabsch & Sander, 1983). The N- and C-terminal capping domains are shown in yellow, the LRR core in blue. A disulphide bond in the N-terminal capping domain is shown in bonds representation, genetic alleles previously characterised for SRF9/SUB are highlighted in magenta). (*e*) Example region of the SRF6 structure covering residues 50-53 and 62-64 (in yellow, in ball-and-stick representation) and including the final (2*F_o_-F_c_*) electron density map contoured at 1.5 σ (blue mesh). (*f*) Structural comparison of the C-terminal capping domains in the Arabidopsis receptor kinases SERK1, SRF6 and SOBIR1. The LRR ectodomain (in blue) structures of SERK1 (r.m.s.d. is ∼ 1.5 Å comparing 163 corresponding C_α_ atoms) and SOBIR1 (r.m.s.d. is ∼ 3.4 Å comparing 133 corresponding C_α_ atoms) were superimposed to the SRF6 LRR domain and presented side-by-side in ChimeraX. The C-terminal capping domains are highlighted in yellow.

**Table 3.**
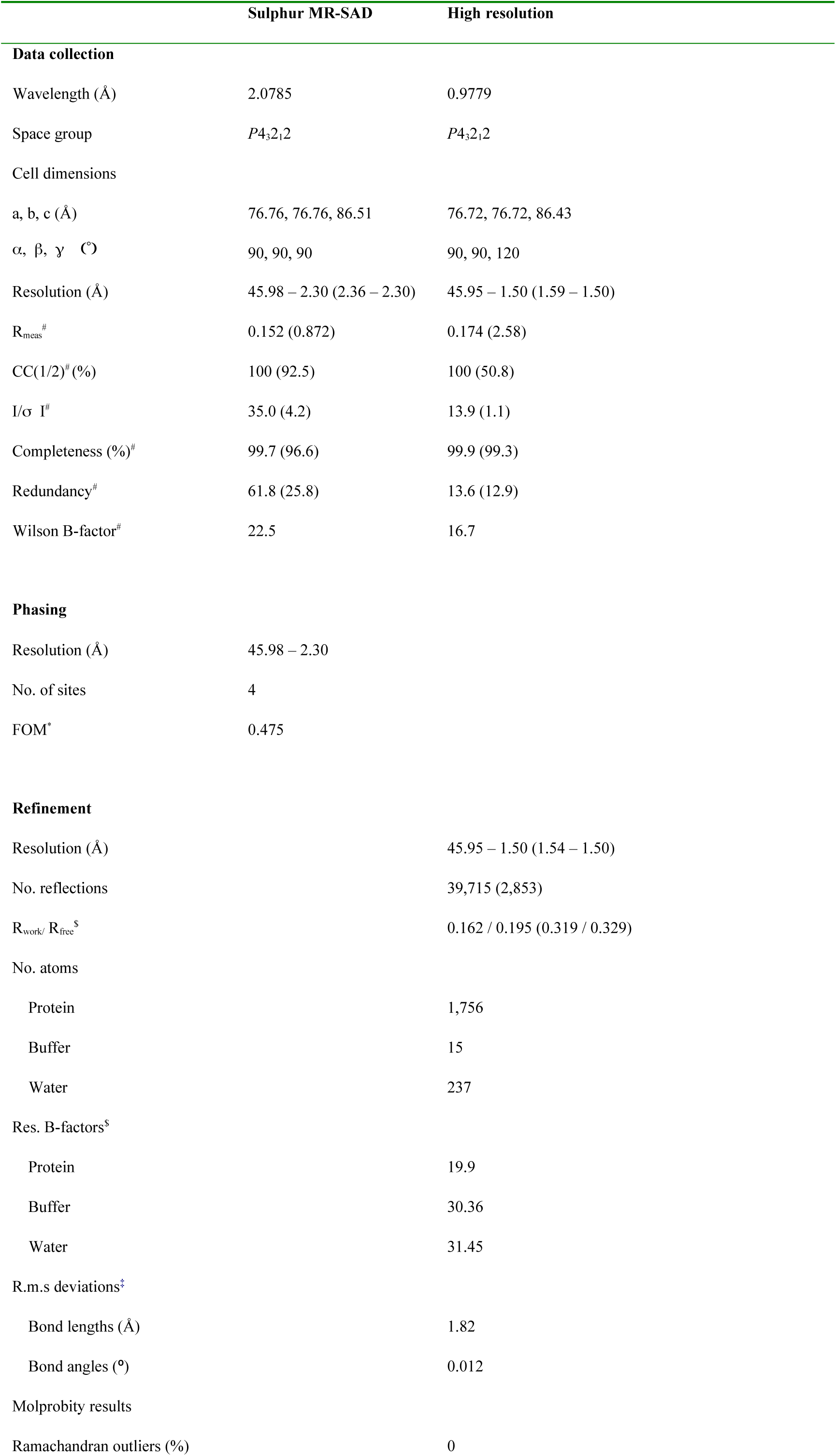
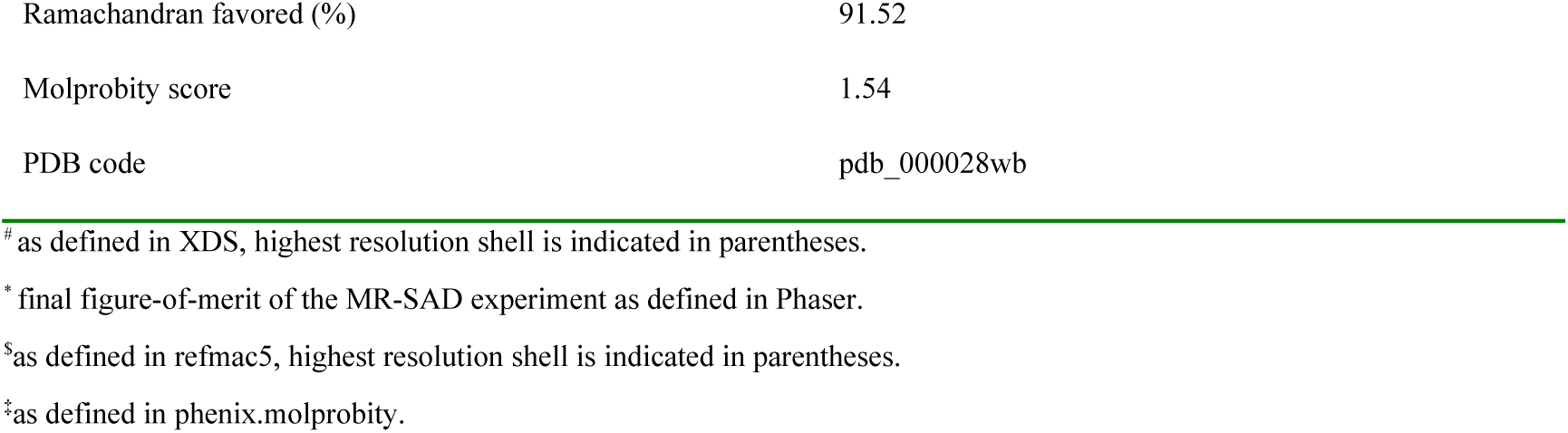
SRF6 crystallographic data collection, phasing and refinement statistics.

### 3.3. Assessing the interaction between the SRF6 and SRF7 ectodomains and BR signalling components

Next, we sought to characterise the potential roles of SRF RKs in brassinosteroid signalling. Previous studies have reported that SRF6 gene expression is upregulated following brassinosteroid treatment (Eyüboglu *et al*., 2007). In addition, high-throughput interaction assays revealed that the ectodomains of SRF4, SRF6, SRF7, and SRF9 interact with the LRR ectodomains of the brassinosteroid receptors BRI1 and BRL1, the SERK co-receptor kinases, and the negative regulators BIR1-4 (Smakowska-Luzan *et al*., 2018). Furthermore, the potato SRF receptor StLRPK1 was reported to constitutively interact with the co-receptor kinase SERK3 in co-immunoprecipitation assays (Wang *et al*., 2018).

We purified the LRR ectodomains of SRF6 and SRF7 (see Methods) and examined their interactions with components of the brassinosteroid signalling pathway. When mixed at equimolar ratios in the absence of brassinosteroids, the BRL1 and SRF7 ectodomains did not co-migrate in analytical size-exclusion chromatography assays (Fig. 3a). Consistently, no interaction was detected by isothermal titration calorimetry (Fig. 3b).

**Figure 3.**
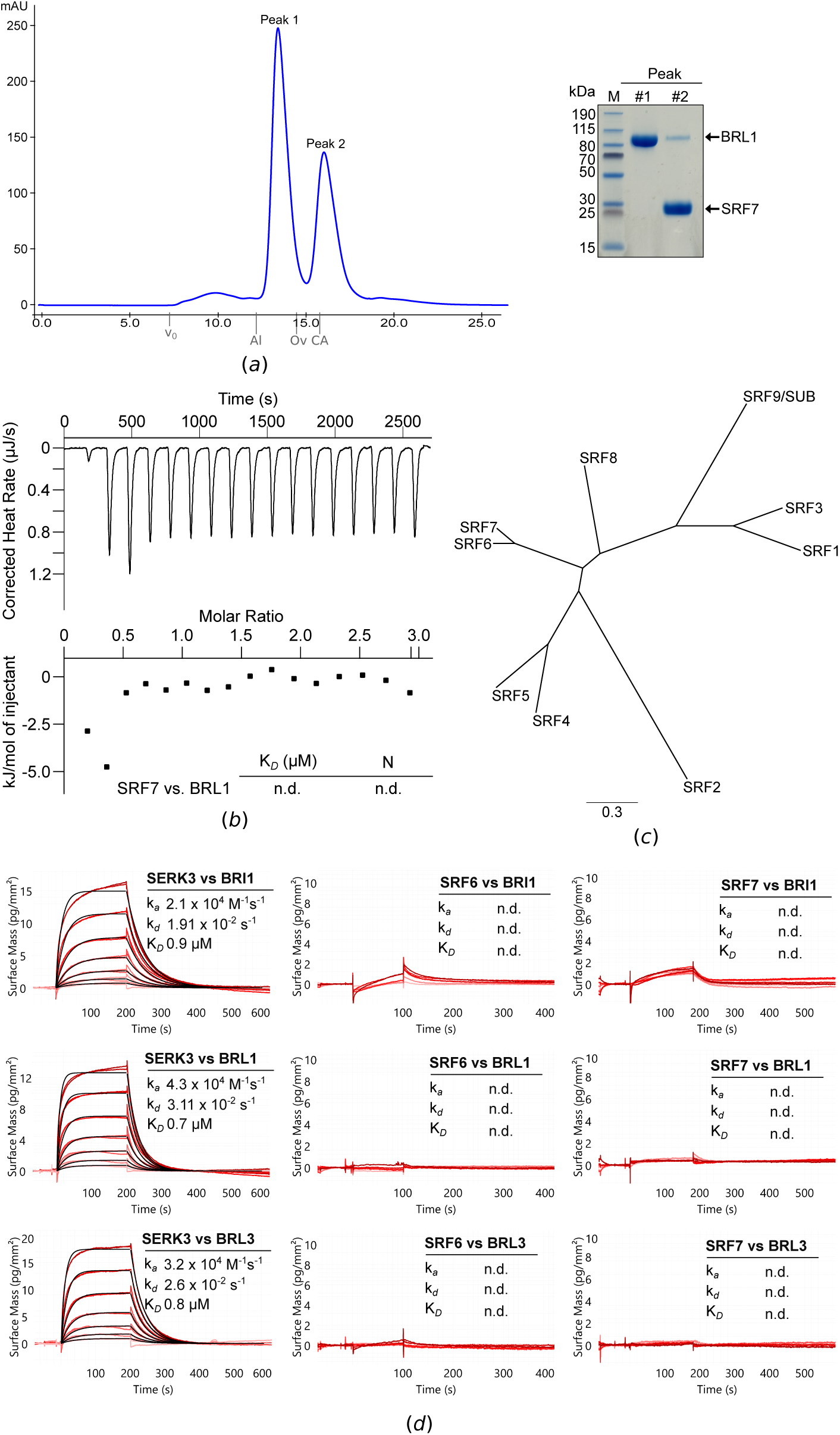
The ectodomains of the SRF6 and SRF7 proteins do not display high-affinity binding to the Arabidopsis brassinosteroid receptors BRI1, BRL1 and BRL3. (*a*) Analytical size-exclusion chromatography of an equimolar mixture of BRL1 and SRF6. The absorbance trace at λ=280 nm is shown in blue. Indicated are the void volume (v_0_) and the elution volumes for molecular-mass standards (Al, Aldolase, 158 kDa; Ov, Ovalbumin, 43 kDa; CA, Carbonic Anhydrase, 29 kDa). A Coomassie-stained SDS-PAGE of the peak fractions is shown alongside. (*b*) Isothermal titration calorimetry of SRF6 (in the syringe) vs. BRL1 (in the cell). Shown are integrated heat peaks (upper panel) versus time and binding isotherms versus molar ratio of SRF6 ligand (lower panel). (*c*) Phylogenetic tree of the 9 SRF family members annotated in the Arabidopsis genome. (*d*) Grating coupled interferometry (GCI) binding kinetics of BRI1, BRL1 and BRL3 vs. SRF6 and SRF7. The known BR co-receptor kinase SERK3 served as positive control. Shown are sensorgrams with raw data in red and their respective fits in black. Binding kinetics were analysed by a 1:1 binding model. Table summaries of kinetic parameters are shown alongside (k*a*, association rate constant; k*d*, dissociation rate constant; K*D*, dissociation constant; n.d., no detectable binding).

As an alternative approach, we analysed the binding of SRF6 and its close homologue SRF7 (Fig. 2 c; ∼70% amino acid sequence identity) by grating-coupled interferometry (GCI). The LRR ectodomains of the brassinosteroid receptor kinases BRI1, BRL1, and BRL3 bound the co-receptor kinase SERK3 in the presence of the steroid ligand brassinolide, consistent with previous reports (Fig. 2d) (Hohmann, Santiago *et al*., 2018; Caregnato *et al*., 2025). In contrast, no binding was detected when either the SRF6 or SRF7 ectodomain was used as the analyte Fig. 3d).

BIR receptor pseudo-kinases have been identified as negative regulators of brassinosteroid signalling (Imkampe *et al*., 2017), a function that depends on their LRR domains interacting with the ectodomains of SERK co-receptor kinases (Ma *et al*., 2017; Hohmann, Nicolet *et al*., 2018). The LRR domains of BIR1-3, immobilised on the GCI chip (see Methods), all interacted with the SERK3 ectodomain, as previously reported (Hohmann, Santiago *et al*., 2018) (Fig. 4). In contrast, no interaction was detected for SRF6 or SRF7 (Fig. 4). Similarly, no binding was observed when testing interactions between SRF6 or SRF7 and SERK3 coupled to the chip (Fig. 4).

**Figure 4.**
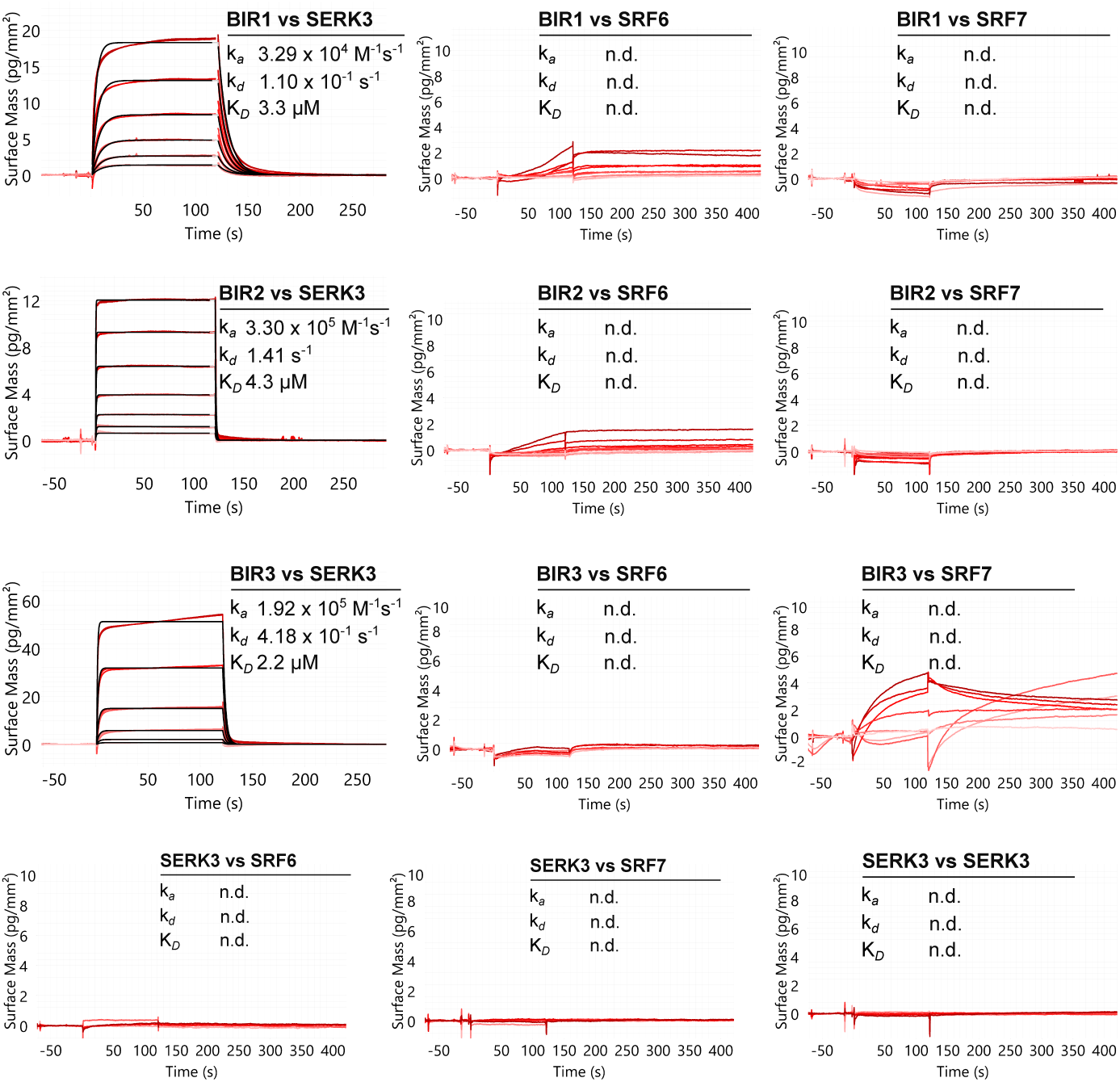
The ectodomains of SRF6 and SRF7 do not display high-affinity binding to Arabidopsis BIR receptor pseudokinases, or to SERK3. Grating coupled interferometry (GCI) binding kinetics of BIR1, BIR2, BIR3 and SERK3 vs. SRF6 and SRF7. The known, ligand-independent interaction between BIRs and SERK3 served as positive control. Shown are sensorgrams with raw data in red and their respective fits in black. Binding kinetics were analysed by a 1:1 binding model. Table summaries of kinetic parameters are shown alongside (k*a*, association rate constant; k*d*, dissociation rate constant; K*D*, dissociation constant; n.d., no detectable binding).

Together, our experiments did not reveal direct, high-affinity interactions between the LRR domains of SRF6 or SRF7 and early components of the brassinosteroid signalling pathway.

## 4. Discussion

Structural biology has provided important insights into the ligand recognition and receptor activation mechanisms of plant membrane receptor kinases (Hohmann *et al*., 2017; Song *et al*., 2017; Moussu & Santiago, 2019). However, the recombinant expression of these proteins and the production of diffraction-quality crystals remain significant challenges. The LRR crystallisation screen described here enabled the successful crystallisation and structure determination of several plant RK ectodomains and ectodomain complexes (Fig. 1). We speculate that the relatively low pH of the screen conditions is a key factor promoting crystallisation of extracellular proteins in otherwise standard PEG- or salt-based crystallisation buffers. The individual conditions can be readily prepared from common crystallisation stock solutions and stored in sterile-filtered 50 ml Falcon tubes, facilitating the long-term preservation and routine use of this in-house screen.

The SRF6 ectodomain crystallised readily in several LRR screen conditions and its structure could be determined at high resolution by MR-SAD. The resulting model reveals a compact LRR ectodomain architecture that overall resembles other plant LRR receptor kinases and co-receptor kinases, supporting the notion that SRF proteins belong to the structurally conserved LRR-RK superfamily (Fig. 2).

Motivated by previous reports linking SRF proteins to brassinosteroid responses, we have examined whether the SRF6 and SRF7 ectodomains interact directly with key components of the brassinosteroid signalling pathway. However, using multiple complementary biochemical approaches, including analytical size-exclusion chromatography, isothermal titration calorimetry, and grating-coupled interferometry, we were unable to detect direct interactions between the ectodomains of SRF6 or SRF7 and brassinosteroid receptors, co-receptor kinases and receptor pseudo-kinases (Fig. 3, 4)

Taken together, these results suggest that the SRF6 and SRF7 ectodomains do not form stable, high-affinity complexes with early components of the brassinosteroid receptor complex under the conditions tested. This observation contrasts with previous high-throughput interaction screens (Smakowska-Luzan *et al*., 2018) and in planta biochemical assays (Wang *et al*., 2018). One possible explanation is that such interactions may be transient, indirect, or mediated by additional factors present in the cellular context but absent from the in vitro binding assays used here. Alternatively, SRF receptors may participate in signalling pathways that intersect with brassinosteroid responses downstream of receptor activation rather than through direct receptor-receptor interactions at the plasma membrane. Future studies will be required to elucidate the physiological functions of SRF receptors in plant organ development and in cell wall sensing. The extracellular SRF LRR domain may also function as a ligand-binding module for a cell wall-derived signalling molecule (Bhasin *et al*., 2025), an intriguing possibility that warrants further investigation.

## Acknowledgements

We thank Y. Belkhadir for discussions and the staff of beamline X06DA (PXIII) at the Swiss Light Source (SLS) Villigen, Switzerland for technical help during data collection. Raw diffraction images and XDS processing files have been deposited with DOI 10.5281/zenodo.18745495.

## PDB reference

Crystal structure of the STRUBBELIG-RECEPTOR FAMILY 6 (SRF6) ectodomain from Arabidopsis thaliana, pdb_000028wb.

